# Polygalacturonase production enhancement by Piriformospora indica from sugar beet pulp under submerged fermentation using surface methodology

**DOI:** 10.1101/2021.11.18.469178

**Authors:** Somayyeh Kiani, Parisa Fathi Rezaei, Sina Jamalzadegan

## Abstract

This study proposed a novel and cost-effective approach to enhance and optimize the polygalacturonase from *P. indica*. In current investigation, the impact of ammonium sulfate, sugar beet pulp (SBP) and glucose as variables on induction of polygalacturonase from *P. indica* was optimized using the central composite design (CCD) of response surface methodology (RSM) under SmF. Additionally, partial polygalacturonase purification and in situ analysis were performed. The optimal reaction conditions, which resulted in the highest enzyme activity were observed as the following conditions: ammonium sulfate (4 g/L), SBP (20 g/L), glucose (60 g/L). Under the optimized condition, the maximum enzyme activity reached to 19.4 U/ml (127 U/mg) which increased by 5.84 times compared to non-optimized conditions. The partial purified polygalacturonase molecular weight was estimated 60 KDa. In line with the bioinformatic analysis, exo-polygalacturonase sequence of *P. indica* showed similarity with *Rhizoctonia solani’s and Thanateporus cucumeris*. These results indicated that SBP act as a cheap and suitable inducer of polygalacturonase production by *P. indica* in a submerged cultivation. The outcome of this study will be useful for industries to decrease environmental pollution with cost-effective approaches.

## 1. Introduction

Pectinases or pectin depolymerases known as a very important industrial enzymes have a broad range of applications and play a crucial role in industrial biotechnology [1]. The pectinolytic enzymes are produced by higher plants, bacteria, fungi, yeasts under submerged and solid-state fermentation conditions. They are classified regarding to their mechanism of action: pectin esterase, pectinase (polygalacturonase) and pectin lyase; and caused to production of galacturonic acid [2]. Like many other depolymerizing enzymes, they are usually inducible by the polymer they degrade. Fungal pectinases as main industrial enzymes and are of great significance with extensive application [3]. The important steps in pectinase upstream process include; the choice of source, substrate, reaction conditions and reactor design [1].

Pectin as the acidic heteropolysaccharide is mainly composed of galacturonic acid which present as the major components of middle lamella and primary cell wall of plants [4]. Nowadays, in considering the global sensitivity about the environment, enzyme production from wastes, caused to overcome the problem of high-cost production in the industry and prevent environmental pollution [5]. With the increasing application of pectinase, decreasing cost production has become one of the most important targets. Previous researches have mentioned that pectin-containing agro-wastes, including sugar beet pulp, citrus pulp pellets, apple pomace, henequen pulp, lemon pulp and other related materials as carbon source could be induced pectinase production by many microorganisms [6]. Sugar beet pulp (SBP) as the by-product of the beet sugar industry, is produced annually in large quantities. On the other hand, SBP could be an important renewable resource and its bio-conversion appears to be of great biotechnological importance. The lignocellulosic portion of dried SBP is consist of 22– 30% cellulose, 24–32% of hemicellulose (essentially arabinan), 24–32% of pectins substances and 3–4% of lignin [7]. Due to high pectin content of SBP, it could be used for pectinolytic enzymes production without adding any pectinaceous materials as enzyme inducer [8].

The induction of pectinase production by various organisms from agricultural by-products were described; by *Penicillium fellutanum* from wheat bran [9], by *Bacillus pumilus* from mixture of banana and orange peel [10], by *Aspergillus niger* DMF 27 and DMF 45 from deseeded sunXower head [3], by *Aspergillus niger* from citrus waste peel [11], by *Aspergillus sojae* from agricultural and agro-industrial residues [12], by *Aspergillus niger* and *Bacillus gibsoni* from sugar beet pulp [13], by *T. reesei* Rut C-30 from sugar beet pulp [14], and by *Bacillus pumilus* from sugar beet pulp and wheat bran [5].

In the current investigation, the production of exo-polygalacturonase from *P. indica* by sugar beet pulp (SBP) as an inducer was optimized by response surface methodology (RSM) and its molecular characteristics evaluated by partial purification and in situ analysis.

## 2. Material and methods

### 2.1. SBP preparation

SBP was prepared from Moghan Agro-Industry & Livestock Co., dried at 60 °C for 24 h, the dried SBP grinded and stored in air tight container. The particles with mesh sieve size adjusted to 500 μm were used for submerged fermentation.

### 2.2. Microorganism, media and culture conditions

The *P. indica* fungus was selected for production of polygalacturonase and obtained from the Department of Plant Pathology, School of Agriculture, Tarbiat Modares University (Iran). The fungi were cultured on modified Kaefer medium [15] and glucose was replaced with SBP. SBP^+^ represents the SBP-containing medium.

For submerged cultivation of *P. indica*, 10 mm of agar discs were transferred to 250 mL flasks containing 50 mL of modified Kaefer medium supplemented with 10 g/L SBP and incubated in shaker (200 rpm) at 29 °C for 10 days. Medium without sugar beet pulp (SBP^-^) was used as control. The samples were assayed for pectinase activity and fungal growth measurement.

### 2.3. Measurement of cell fresh and dry weight, growth yield and specific growth rate

At the end of each incubation time, the culture broth was filtered through Whatman No. 1 paper and growth parameters including; growth yield (Y_X/S_), specific growth rate (μ) and spore yield fresh and medium pH were determined [16].

### 2.4. Total protein determination

Protein content and pectinase activity were determined in the cell-free supernatant after centrifugation of culture broth at 12880 rcf at 4 °C for 15 min. Determination of total protein content was performed according to the Bradford’s method and bovine serum albumin used as the standard [17].

### 2.5. Pectinase activity

Polygalacturonase activity was evaluated by measuring the released reducing end products, using 3, 5-dinitrosalicylic acid (DNS) and expressed as galacturonic acid equivalent [18]. The enzymatic reaction mixture included 0.25 ml of cell-free supernatant and 0.75 ml of 1% pectin in 0.2 M phosphate buffer pH 6.5 as substrate. The mixture was incubated at 60 °C for 5 min. One unit (U) was expressed in term of the enzyme quantity which would yield 1μmol galacturonic acid per minute during the standard assay condition.

### 2.6. Identification of the significant variables using experimental design

In order to maximize enzyme production and understand the role of interacting variables, optimization of the medium constituents was done by central composite design (CCD). Three variables including; glucose (A), ammonium sulfate (B) and SBP (C) were selected to find the optimized condition for the production of pectinase and twenty experimental runs with three center points generated including the response surface plot by using the statistical software package Design-Expert 7.0.0 (Stat Ease Inc., Minneapolis, USA). The range and the levels of the variables are given in Table 1. The recommended 20 experiments by using different composition of independent variables was shown in Table 2. Statistical analysis of the model was performed to evaluate the analysis of variance (ANOVA). The quality of the polynomial model equation was judged statistically by determination coefficient R^2^, and its statistical significance was determined by F-test.

**Table 1.**
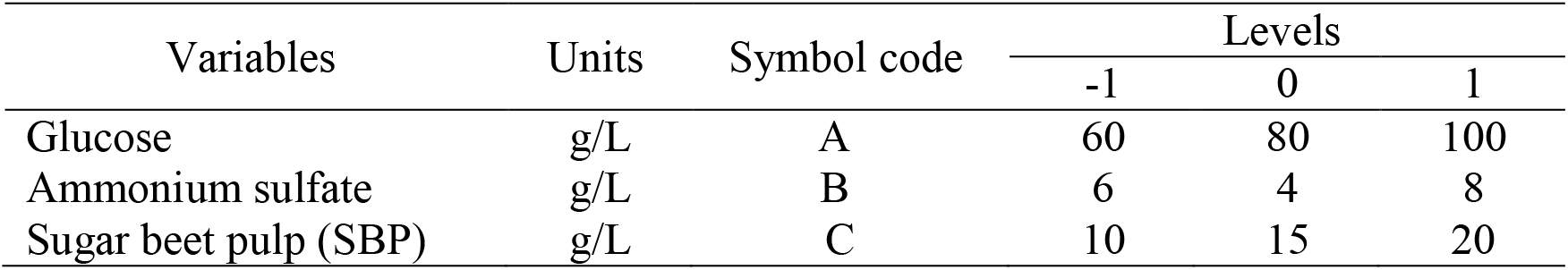
Levels of independent variables used in CCD design.

| Variables | Units | Symbol code | Levels |  |  |
| --- | --- | --- | --- | --- | --- |
|  |  |  | -1 | 0 | 1 |
| Glucose | g/L | A | 60 | 80 | 100 |
| Ammonium sulfate | g/L | B | 6 | 4 | 8 |
| Sugar beet pulp (SBP) | g/L | C | 10 | 15 | 20 |

**Table 2.**
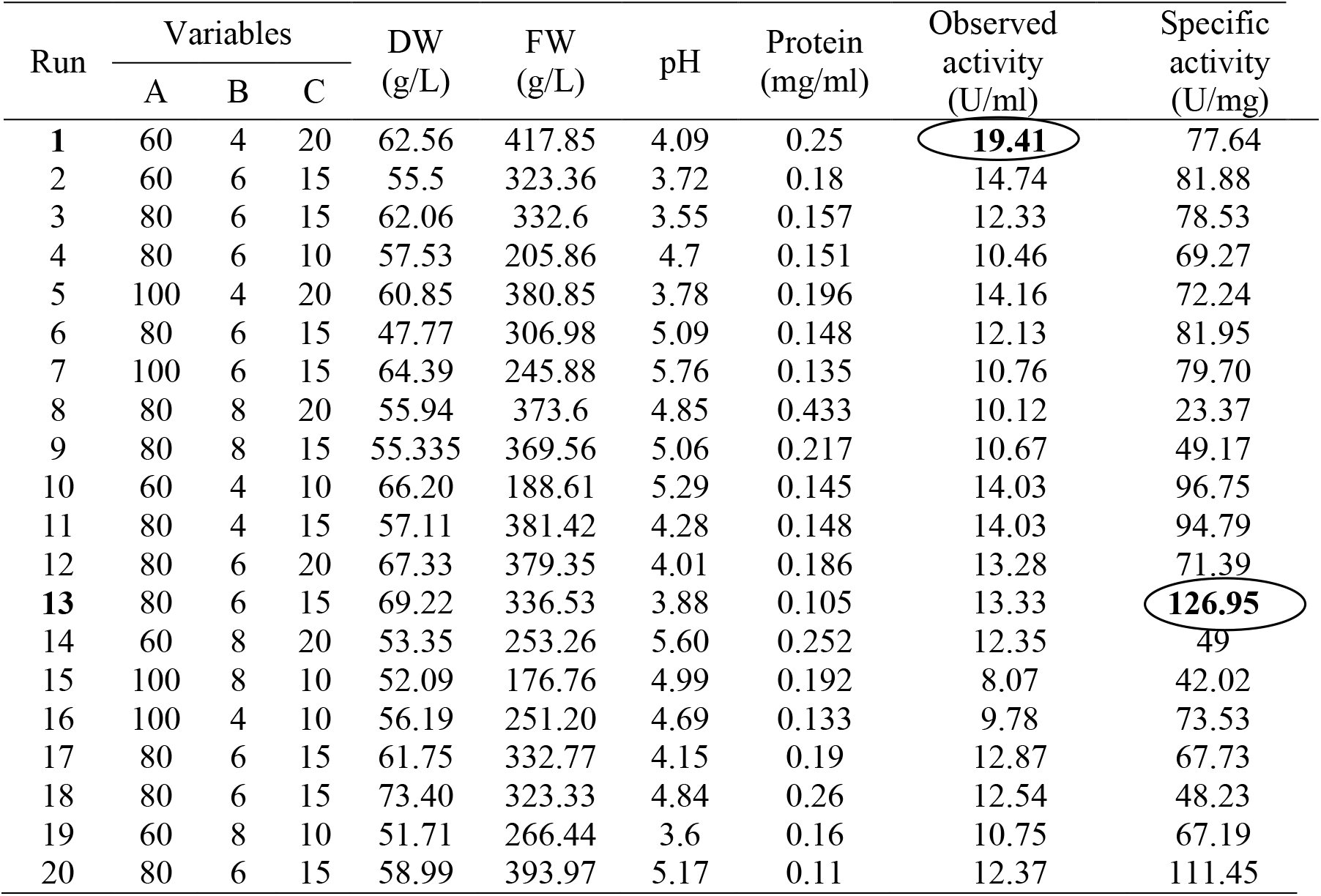
CCD design matrix for pectinase production factors and corresponding results.

### 2.7. Pectinase partial purification

To purify the pectinase, the fungus was cultured in 250 mL shake flasks with 100 mL of optimized fermentation medium. The fermentation broth was separated by centrifugation at 10,000 g for 15 min at 4°C and the cell-free supernatant was saturated with ammonium sulphate to 90% saturation. The saturated solution was left overnight at 4ºC with gentle agitation, centrifuged at 10000 g for 20 min at 4 ºC, the precipitate solubilized in minimal amount of 10 mM sodium acetate buffer (pH 5.75) and dialyzed against the same buffer for 24 h at 4ºC. The obtained-dialyzed proteins were used for enzyme characterization. The protein content and enzyme activity were determined as described in earlier part.

### 2.8. Gel electrophoresis

The molecular mass and purity of polygalacturonase were determined by SDS-PAGE (12.5% running gel and 5% stacking gel) Laemmli (1970) [19]. The protein samples were denatured by heating at 100°C with the sample buffer for 5 min before loading and the gel was stained by silver staining method of Merril et al [20].

### 2.9. Phylogenetic tree simulation

We applied Molecular Evolutionary Genetics Analysis (MEGA X) software as a powerful tool for constructing sequence alignments, gathering phylogenetic histories, and performing molecular evolutionary analysis. This software can be used for comparing DNA and protein sequences. Firstly, we aligned the DNA sequences of more than fifty different strain types of extracellular polygalacturonases and then a phylogenetic tree was constructed for those data by the maximum likelihood method. In this approach, an initial phylogenetic tree was constructed using a Neighbor-Joining, and its branch lengths are modified to maximize the likelihood of the data set for that tree topology under the desired model of evolution. Then the NNI (nearest neighbor Interchange) approach was used for creating the variants of the topology. NNI tries to search for topologies that are in good shape with the data better. The search is repeated until no greater likelihoods are found. Finally, the Neighbor-joining tree of different extracellular polygalacturonase strains was constructed after 500 iterations and the bootstrap confidence values were calculated and shown on node in Fig. 4. The protein sequences were gathered from Uniprot and GenBank (NCBI) [21] [22] (https://academic.oup.com/nar/article/22/22/4673/2400290?login=true). [23]

## 3. Results

### 3.1 Growth of fungus

Production of pectinase by *P. indica* was evaluated for 10 days (Fig. 1). The production of pectinase on both medium reached maximum on 6th day of culture and then decreased. The production of enzyme on SBP^−^ and SBP^+^ medium was determined 2.2 and 3.32 U/ml, respectively. Also, the highest amount of dry and fresh weight on both medium was detected on 6th day of culture (Fig. 1). As shown in Table 3, the lowest and highest dry cell weight were detected on unmodified Kaefer medium and SBP and glucose containing medium, respectively. Furthermore, the highest amount of growth yield and specific growth rate were measured on medium containing ammonium sulfate, glucose and SBP, 0.62 and 1.61 respectively.

**Table 3.**
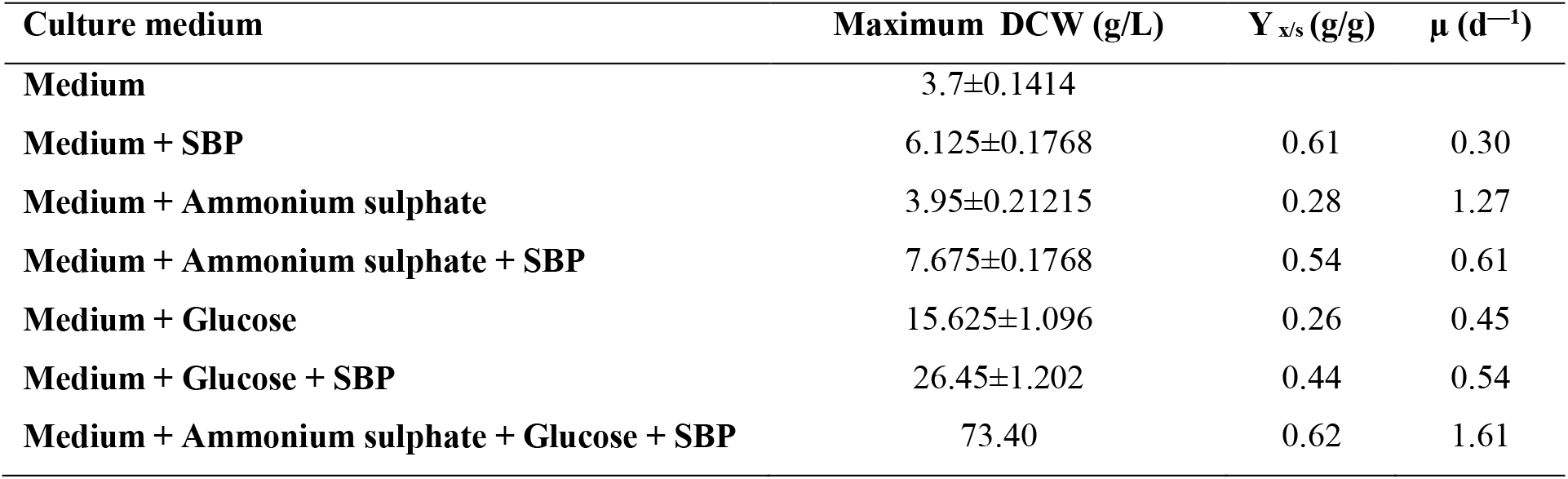
Effect of variables on growth of *P. indica*.

| Culture medium | Maximum DCW (g/L) | $Y_{x/s}$ (g/g) | $\mu$ (d<sup>-1</sup>) |
| --- | --- | --- | --- |
| Medium | 3.7±0.1414 |  |  |
| Medium + SBP | 6.125±0.1768 | 0.61 | 0.30 |
| Medium + Ammonium sulphate | 3.95±0.21215 | 0.28 | 1.27 |
| Medium + Ammonium sulphate + SBP | 7.675±0.1768 | 0.54 | 0.61 |
| Medium + Glucose | 15.625±1.096 | 0.26 | 0.45 |
| Medium + Glucose + SBP | 26.45±1.202 | 0.44 | 0.54 |
| Medium + Ammonium sulphate + Glucose + SBP | 73.40 | 0.62 | 1.61 |

**Fig. 1.**
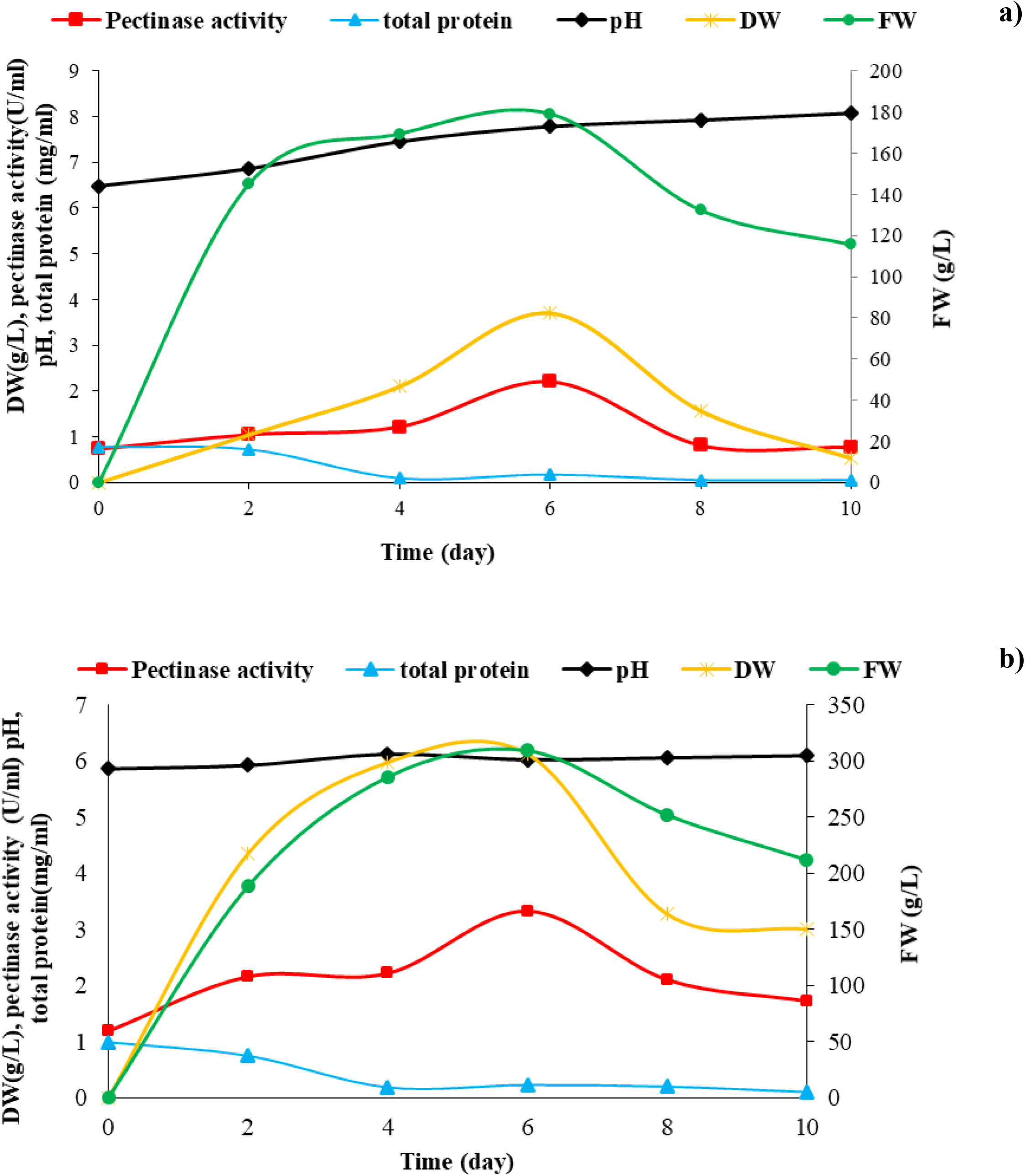
The effect of SBP on growth parameters and pectinase activity of *P. indica*. Time course profile of *P. indica* on Kaefer medium (a) and supplemented with SBP (b) in SmF. Data are shown as mean ± SD of three independent experiments in triplicate layout. SBP^+^: medium containing SBP, SBP^-^: medium without SBP, FW: Fresh weight; DW: dry weight.

### 3.2 Optimization of the pectinase production by RSM

Three variables that have the maximum effect on the polygalacturonase production were determined by one-factor-at-a-time method and the interaction between various selected factors on polygalacturonase production (glucose, ammonium sulfate and SBP concentration) were investigated by RSM.

Then the results were analyzed by standard analysis of variance (ANOVA) and the CCD design was fitted with the second-order polynomial equation:

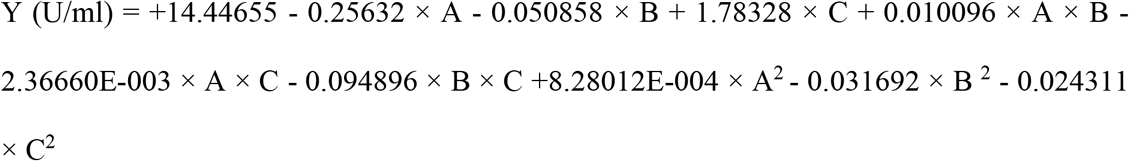

Eq. (1) polygalacturonase activity (Y) as a function of Glucose (A), Ammonium sulfate (B) and Sugar beet pulp (C).

The software suggested 20 experiments and the predicted and experimental values for enzyme production are presented in Table 2. The sufficiency of the model was checked using correlation coefficient (R^2^) and the closer the value of R^2^ to 1, the better the correlation between the observed and the predicted values. The correlation coefficient (R^2^) which shows the relationship between the experimental and predicted responses was 0.9866 and thus the model could explain more than 98.66% of the variability in the responses (Table 4).

**Table 4.** ANOVA for response surface quadratic model of polygalacturonase production.

| Source | Sum of squares | df | Mean squares | F value | P – value |  |
| --- | --- | --- | --- | --- | --- | --- |
| Model | 106.59 | 9 | 11.84 | 81.76 | <0.0001 | significant |
| A | 29.23 | 1 | 29.23 | 201.82 | <0.0001 |  |
| B | 38.49 | 1 | 38.49 | 265.70 | <0.0001 |  |
| C | 19.13 | 1 | 19.13 | 132.08 | <0.0001 |  |
| AB | 0.92 | 1 | 0.92 | 6.35 | 0.0304 |  |
| AC | 0.32 | 1 | 0.32 | 2.18 | 0.1705 |  |
| BC | 6.14 | 1 | 6.14 | 42.37 | < 0.0001 |  |
| A<sup>2</sup> | 0.31 | 1 | 0.31 | 2.15 | 0.1733 |  |
| B<sup>2</sup> | 0.047 | 1 | 0.047 | 0.32 | 0.5829 |  |
| C<sup>2</sup> | 1.07 | 1 | 1.07 | 7.40 | 0.0215 |  |
| Residual | 1.45 | 10 | 0.14 | - | - |  |
| Lake of Fit | 0.50 | 5 | 0.10 | 0.53 | 0.7483 | Not significant |
| Pure error | 0.95 | 5 | 0.19 |  |  |  |
| Cor total | 108.04 | 19 | - | - | - |  |

Note: R<sup>2</sup> (0.9866), Adj R<sup>2</sup> (0.9745), Pred R<sup>2</sup> (0.9404), Adeq precision 41.209.
P-values < 0.05 indicate significant differences.

Moreover, R^2^ values were in reasonable agreement with adjusted R^2^ values of 0.9745 (polygalacturonase production). Values greater than 0.1000 indicate the model terms are not significant. The “Pred R-Squared” of 0.9404 is in reasonable agreement with the “Adj R-Squared” of 0.9745. Table 4 represents the obtained results of the quadratic response surface model fitting in the form of ANOVA.

The model F-value is 81.76 which indicates the model significance. The (B) had the highest F-value of 265 implying that it had the most significant influence on the enzyme activity in comparison to glucose (A) and SBP (C). Moreover, the lack of fit F-value was 0.53 which is nonsignificant relative to the pure error. The model is geared toward perfect fitness.

### 3.3 Interaction between operating factors

According to the ANOVA Table 4, the significancy of the independent variables and interaction between them was determined by F-values and p-values. The lower the p- value implies the greater the statistical significance of the observed difference. The greater value of F-value illustrates that the factors appropriately estimate the variation of the data about its mean, and the selected factor effects are actual.

As seen in Table 4, A, B, C, AB (glucose and Ammonium sulfate) and BC (Ammonium sulfate and SBP) with a very small p-value (p<0.05) were significant where as, AC was insignificant on enzyme production.

Due to the positive linear coefficient of SBP, by increasing the SBP concentration within the range assayed increase enzyme production and the negative quadratic coefficients of ammonium sulfate and SBP explain the maximum enzyme production at these levels and the enzyme production decreased out of this point. In accordance with the coefficients, SBP is determined as the factor with the most positive significant impact on enzyme production.

Furthermore the interaction among variables was confirmed by the 3-D response surface plots which is used to identify the optimum levels and the interaction between variables evaluated in pectinase production.

The 3-D plots represent the interaction between the two factors, while the other factor was fixed at its optimum level for maximum enzyme production (Fig. 2). Fig. 2a exhibits the interaction between ammonium sulfate and glucose, reveals that pectinase production increases by increasing in ammonium sulfate and glucose concentrations. The response between glucose and SBP indicated that increasing the SBP content and decreasing glucose concentration led to increase in enzyme production (Fig. 2.b). The plot for the interaction between SBP and ammonium sulfate (Fig. 2c) showing an increase in enzyme production at low level of ammonium sulfate and high level of SBP.

**Fig. 2.**
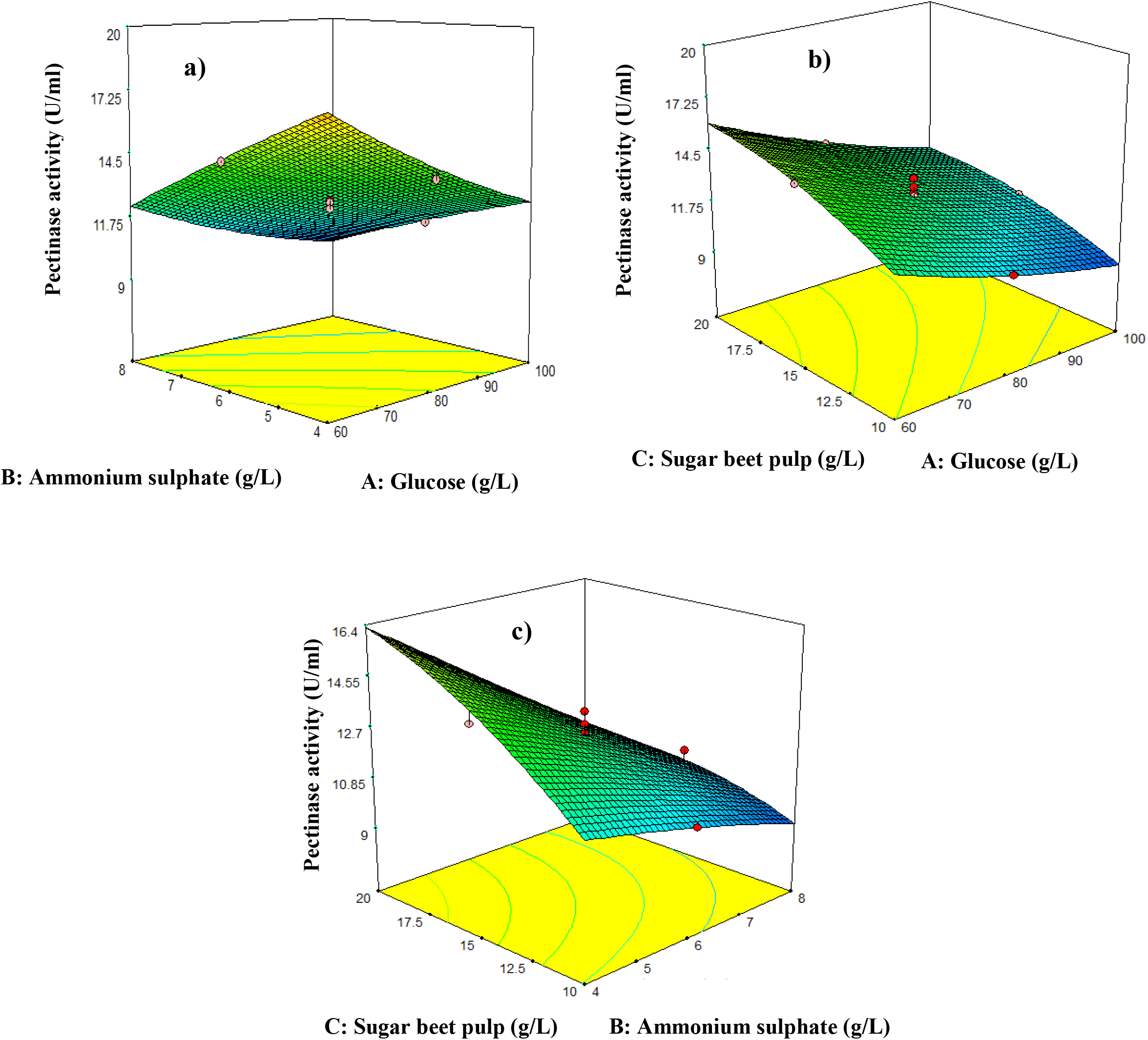
Response surface plot of polygalacturonase yield under optimal conditions and interaction between variables. a) interaction between ammonium sulfate and glucose; b) interaction between sugar beet pulp and glucose; and c) interaction between sugar beet pulp and ammonium sulfate.

The maximum enzyme production was occurred at low levels of glucose and ammonium sulfate and high levels of SBP.

Effect of 3 variables on the response were significant (P > 0.05) which showed high contribution to the enzyme production in SmF (Table 4).

Finally, the optimum condition for pectinase production obtained at 60 g/L of glucose, 4 g/L of ammonium sulfate and 20 g/L of SBP. At the optimum condition, the enzyme activity increase to 19.41 U/ml which is 5.58 fold more than unoptimized condition.

### 3.4 Partial purification of pectinase

For purification, the supernatant was precipitated by ammonium sulfate. The partial purification results were shown in Table 5. Then the specific activity and fold purification was calculated 63.77 and 0.82 respectively. The molecular mass of pectinase was found around 60 kDa by SDS-PAGE (Fig. 3).

**Table 5.** Partial purification process of pectinase.

| Purification step | activity<br>(U/ml) | protein<br>(mg/ml) | Specific activity<br>(U/mg) | Recovery<br>(%) | Purification fold |
| --- | --- | --- | --- | --- | --- |
| Crude extraction | 19.40 | 0.25 | 77.6 | 100 | 1 |
| Ammonium sulfate<br>precipitation | 11.48 | 0.18 | 63.77 | 59.05 | 0.82 |

**Fig.3.**
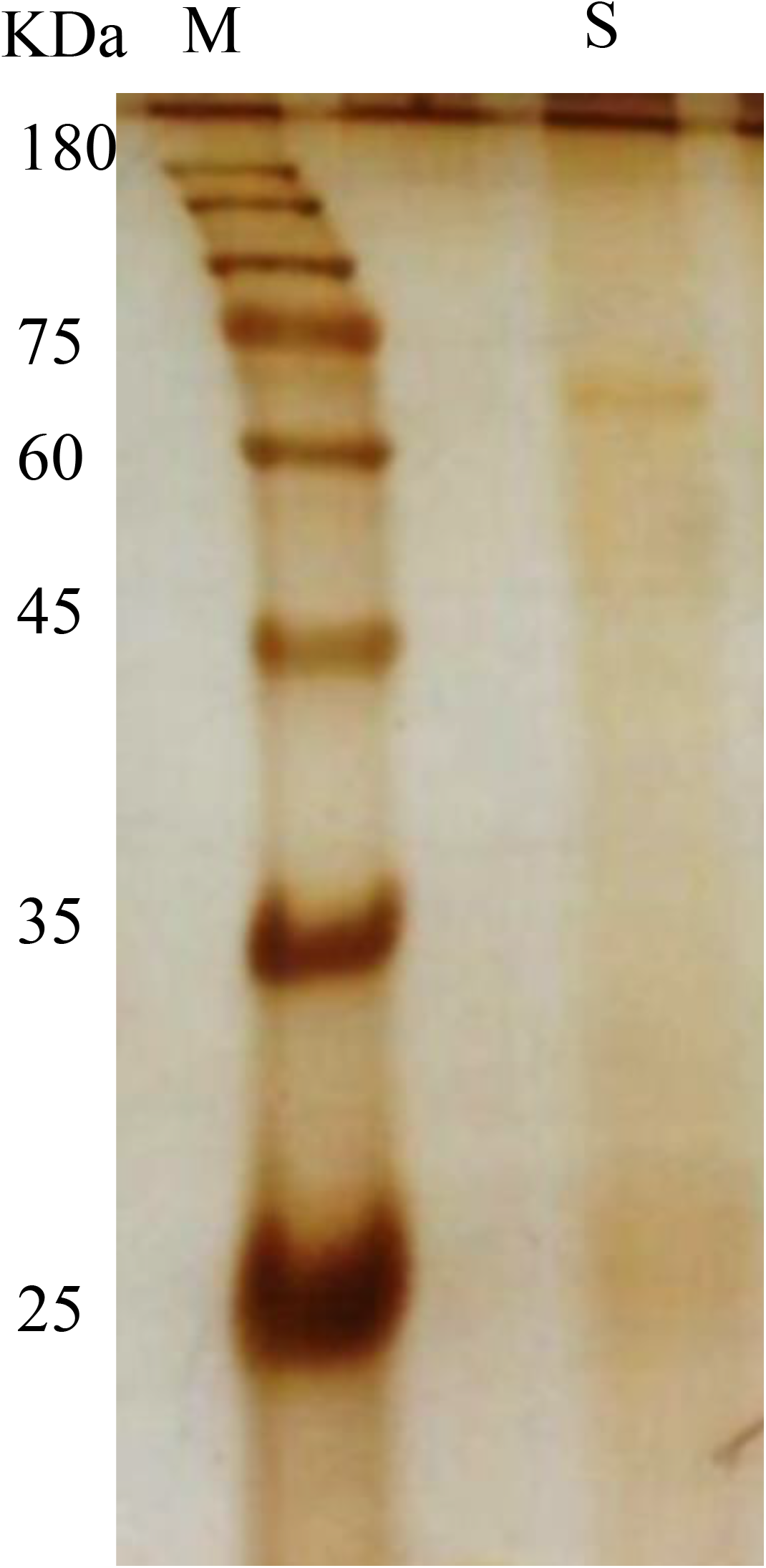
SDS-PAGE analysis of polygalacturonase. M: protein molecular weight marker; S: the sample after partial purification.

### 3.5 Simulation results

As shown in Fig.4, the phylogenetic tree gathered from the concatenated rDNA and TEF alignment by applying heuristic ML analysis with the bootstrap values are shown on nodes. We compared the phylogenetic relationships of *P. indica* with *Rhizoctonia solani’s* extracellular polygalacturonases in NCBI data bank. In Fig.5, *Rhizoctonia solani’s and Thanateporus cucumeris* revealed exo-polygalacturonases similar sequences. Also in Fig. 6 we showed the alignment of the predicted amino acids sequences of *P. indica* polygalacturonase with 8 similar sequences:

**Fig.4.**
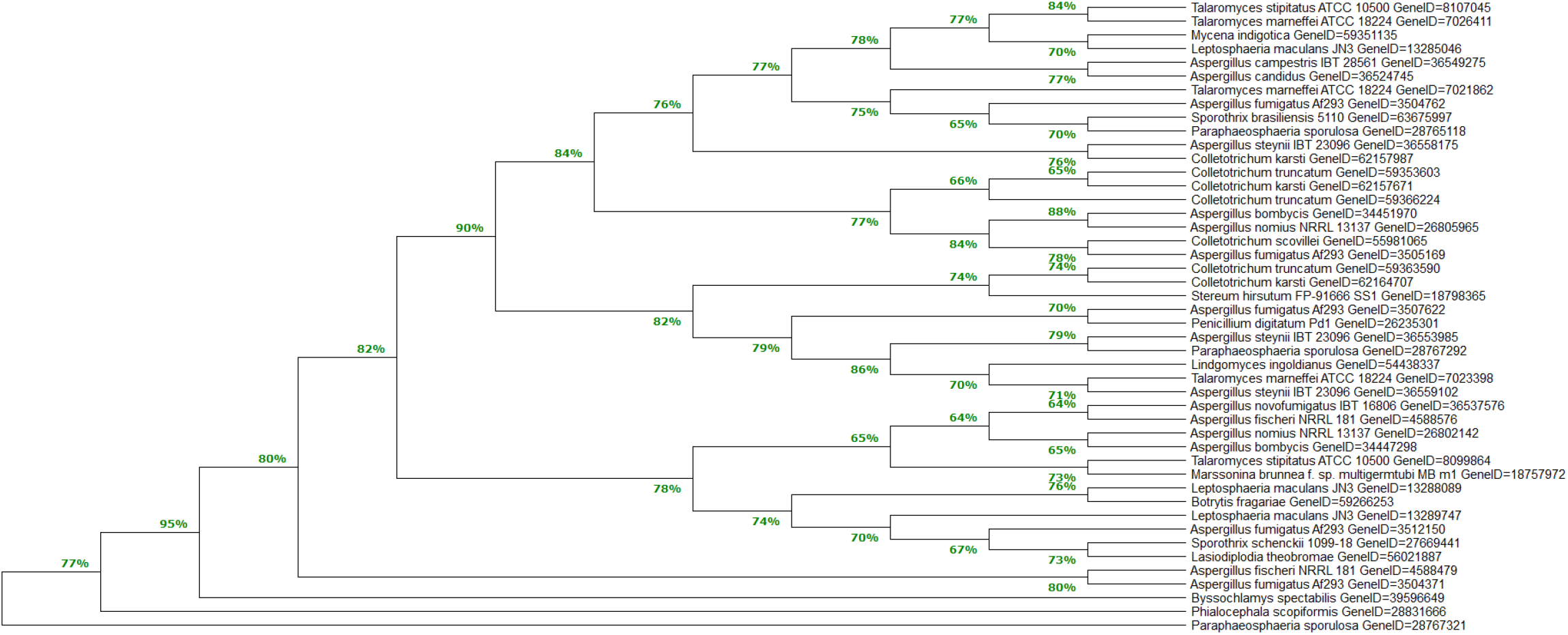
Phylogenetic tree of the *P. indica* and closely related proteins created using the neighbor-joining method. Bootstrap confidence values (500 repetitions) are shown on nodes. The numbers at each node marked the percentage of supporting bootstrap samples.

**Fig. 5.**
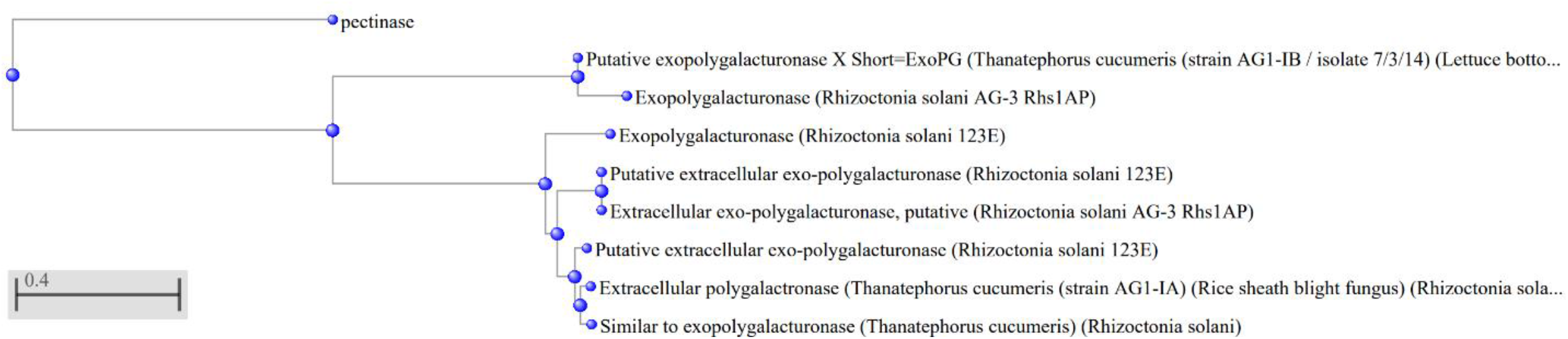
The phylogenetic tree represents similar DNA sequences to the polygalacturonase gene of *P. indica*. The protein sequences were retrieved from Uniprot and GenBank (NCBI).

**Fig. 6.**
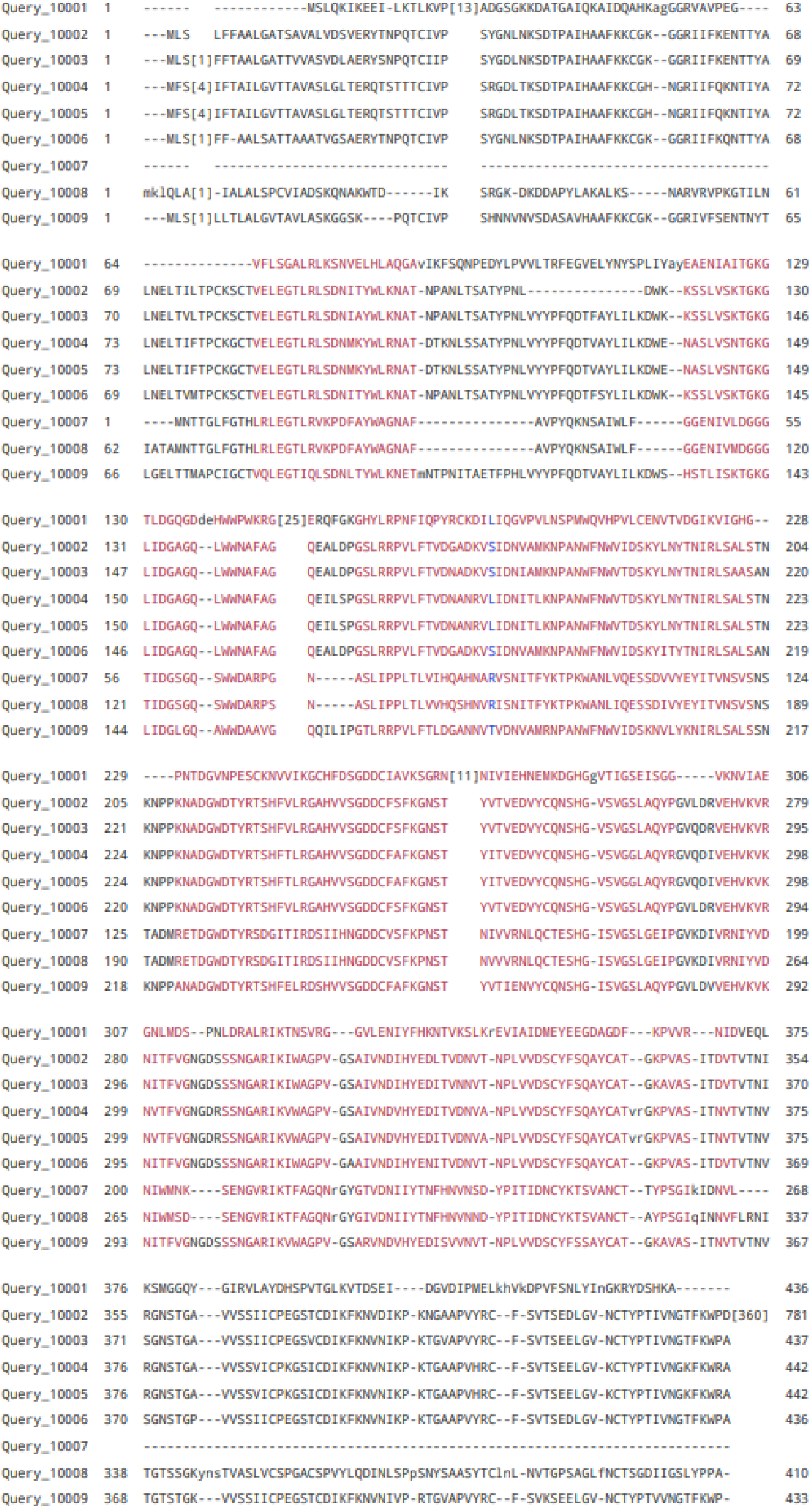
Multiple alignment of the amino acid sequence of *P. indica* polygalacturonase and *Rhizoctonia solani*. The dashes exhibit gaps to improve the alignment. Conserved residues, no gaps, gaps are represented in red, blue and gray, respectively. Query_10001: pectinase, Query_10002: Extracellular polygalacturonase (*Thanatephorus cucumeris* (strain AG1-IA) (Rice sheath blight fungus) (*Rhizoctonia solani*)), Query_10003: Putative extracellular exo-polygalacturonase (*Rhizoctonia solani* 123E), Query_10004: Putative extracellular exo-polygalacturonase (*Rhizoctonia solani* 123E), Query_10005: Extracellular exo-polygalacturonase, putative (*Rhizoctonia solani* AG-3 Rhs1AP), Query_10006: Similar to exo-polygalacturonase (*Thanatephorus cucumeris*) (*Rhizoctonia solani*), Query_10007: Putative exo-polygalacturonase X Short=ExoPG (*Thanatephorus cucumeris* (strain AG1-IB / isolate 7/3/14) (Lettuce bottom rot fungus) (*Rhizoctonia solani*), Query_10008: Exopolygalacturonase (*Rhizoctonia solani* AG-3 Rhs1AP), Query_10009: Exo-polygalacturonase (*Rhizoctonia solani* 123E).

## 4. Discussion

Nowadays, there is a great global interest to obtain pectinase from cheap and suitable substrates like agricultural residues. To evaluate the efficiency of *P. indica* in biodegradation of SBP, the polygalacturonase activity on SBP^+^ and SBP^−^ media have been reported for 10 days (Fig. 1). The polygalacturonase production and fungal growth reached the maximum level simultaneously on both medium. It’s stablished that the components of the medium have great effect on the induction of pectolytic enzymes and several substances play an inducer role on these enzymes synthesis [24]. The utilization of agro-industrial residues, like apple pomace and sugar beet pulp, could notify other substrates and solve environmental problems of the by-products [12]. The choice of perfect agricultural residue to induce enzyme production depends on several factors, such as cost and availability of the substrate material. It is known that duration of fermentation depends on the medium composition, organism, concentration of nutrients and the physiological conditions [3].

SBP has been used as raw material to induce pectinase production by *Aspergillus niger*. It has also been used as the carbon source as well as the pectinase inducer to produce extracellular alkaline pectinase, by *Bacillus gibsoni*, under SSF [13].

The highest activity of exo-pectinase in medium containing sugar beet pulp was determined 3.4 U/mL on 96 h of fermentation and following optimization it reaches to 19.41 U/ml.

Cultivation of *T. reesei* Rut C-30 on sugar beet pulp (50 g/L), the protein content, pectinase activity and specific activity reached their maximum value after 60 h of fermentation (0.43 g/l, 0.82 U/ml and 1.9 U/mg respectively) [14]. The highest enzyme activity by *A. sojae* on 30% sugar beet pulp as an inducer and wheat bran as medium wetted agent attained after 8 days [12].

The greatest endo- and exo-pectinase activity by *A. niger* from sunflower head in SSF (5.1U/g and 17.1U/g) and SmF (4.5 U/ml and 16.0 U/ml) were measured on 96 h. According to the different studies about the fermentation time, it exhibited wide range 40–120 h and 90– 120 h in submerged and solid state fermentations, respectively [3].

The *Colletotrichum* isolated from Argentinian soybean, yielded high amount of the PG (1.08 U/ml) after 7–10 days of incubation and coincide with maximum growth. In medium involving glucose as a sole carbon source decreased polygalacturonase production was monitored [25].

Environmental and nutritional factors are known as two essential factors which affect enzyme production by microorganisms. The pectin and polygalacturonic acid applied as only carbon source of the medium induced synthesis of pectinolytic enzymes by *A. niger* and there was no pectolytic activity in medium containing glucose as an only carbon source. Production of pectin degrading enzymes in the presence of pectin and high glucose concentrations were inhibited although glucose in low concentrations promoted their production. The observed low pectinolytic activity in media with high glucose concentrations is possibly due to provide growth need of organism by the glucose consumption and caused to decrease the pectin lysis. Furthermore, at low glucose concentrations, high pectolytic activity were observed [26]. Our results are in agreement with Fawole et al, which the highest pectinolytic activity was attained at the lowest glucose concentration (60 g/L).

Pectinolytic activity by *A. niger* on medium containing pectin, poly galacturonic acid and glucose at 30°C for 5 days was 17.2, 13.8 and 0 U/ml, respectively [26].

Aguilar and Huitron (1987) showed that high exogenous glucose and galacturonic acid could be influenced endo-PG enzyme production by catabolite repression, whereas glucose had no effect on the exo-PG. The Glucose concentration above 10% (w/w) in the SSF, decreased noticeably the activity of endo and exo-PG [27].

In Solis-Pereyra et al study, exo-PG/gdm and endo-PG/gdm activity by *A. niger* on medium containing 16% (w/w) citric pectin, were 281 U and 152 U. Moreover, inhibited enzyme production and growth were detected on 20- 30% (w/w) pectin concentration [28].

Ammonium sulphate was introduced as the favourable nitrogen source for pectinase production by *A. niger* [26]. Our results are in concurrence with the observations of Sapunova who also stablished that ammonium salts act as stimulator of the pectinase. It has been described that nitrogen limitation decreases the production of polygalacturonase [29].

Bai and *et al* examined the impact of different nitrogen sources on pectinase induction and great enzyme activity measured with ammonium sulfate, yeast extract, soya peptone, soya pulp and MGW[6].

Patil and *et al* examined the impact of ammonium phosphate and sulphate on pectinase production by *A. niger* from sunflower head in both SSF and SmF. They described that ammonium phosphate and sulphate had positive effect on production of pectinase in both fermentation conditions but the increase was very less with ammonium phosphate in comparison to ammonium sulphate. The maximum production of endo-pectinase and exo-pectinase by DMF 27 were recorded in SmF condition 18.9 U/ml and 30.3 U/ml respectively [3].

As stated in many studies the average molecular mass of polygalacturonase are in the range of 35-80 KDa [1]. Different microbial species produced different molecular mass of pectinase enzyme and the difference is as a result of the substrate, nature of microorganism, host cell wall and analytical methods (Oyede, 1998). The molecular weight of *P. indica* polygalacturonase was comparable with previous reports.

## 5. Conclusion

The present study showed that optimized conditions by RSM, yielded high polygalacturonase activity by *P. indica* and SBP was established as a significant enzyme inducer substrate. In situ analysis confirmed the similarity of exopolygalacturonse of *P. indica* with *R. solani*’s enzyme. Application of agricultural and agro-industrial wastes is attractive for enzyme production, economically is valuable and cause to decrease environmental pollution. The outcome of the proposed study will open future pathways for using waste raw materials to produce valuable products with cost-effective and eco-friendly approaches.

This research did not receive any specific grant from funding agencies in the public, commercial, or not-for-profit sectors.

## Supporting information

graphical abstract

## Reference

[1] J. John, K.S. Kaimal, M.L. Smith, P.K. Rahman, P.V. Chellam, Advances in upstream and downstream strategies of pectinase bioprocessing: A review, International Journal of Biological Macromolecules 162 (2020) 1086–1099.

[2] M.K. Patidar, S. Nighojkar, A. Kumar, A. Nighojkar, Pectinolytic enzymes-solid state fermentation, assay methods and applications in fruit juice industries: a review, 3 Biotech 8(4) (2018) 1–24.

[3] S.R. Patil, A. Dayanand, Production of pectinase from deseeded sunflower head by Aspergillus niger in submerged and solid-state conditions, Bioresource technology 97(16) (2006) 2054–2058.

[4] S. Satapathy, J.R. Rout, R.G. Kerry, H. Thatoi, S.L. Sahoo, Biochemical prospects of various microbial pectinase and pectin: an approachable concept in pharmaceutical bioprocessing, Frontiers in Nutrition 7 (2020) 117.

[5] O. Tepe, A.Y. Dursun, Exo-pectinase production by Bacillus pumilus using different agricultural wastes and optimizing of medium components using response surface methodology, Environmental Science and Pollution Research 21(16) (2014) 9911–9920.

[6] Z. Bai, H. Zhang, H. Qi, X. Peng, B. Li, Pectinase production by Aspergillus niger using wastewater in solid state fermentation for eliciting plant disease resistance, Bioresource Technology 95(1) (2004) 49–52.

[7] M. Hutnan, M. Drtil, L. Mrafkova, Anaerobic biodegradation of sugar beet pulp, Biodegradation 11(4) (2000) 203–211.

[8] P.S.-N. Nigam, A. Pandey, Biotechnology for agro-industrial residues utilisation: utilisation of agro-residues, Springer Science & Business Media2009.

[9] F. Amin, T. Arooj, Z.-i.-H. Nazli, H.N. Bhatti, M. Bilal, Exo-polygalacturonase production from agro-waste by Penicillium fellutanum and insight into thermodynamic, kinetic, and fruit juice clarification, Biomass Conversion and Biorefinery (2021) 1–11.

[10] P. Viayaraghavan, S. Jeba Kumar, M. Valan Arasu, N.A. Al-Dhabi, Simultaneous production of commercial enzymes using agro industrial residues by statistical approach, Journal of the Science of Food and Agriculture 99(6) (2019) 2685–2696.

[11] I. Ahmed, M.A. Zia, M.A. Hussain, Z. Akram, M.T. Naveed, A. Nowrouzi, Bioprocessing of citrus waste peel for induced pectinase production by Aspergillus niger; its purification and characterization, Journal of Radiation Research and Applied Sciences 9(2) (2016) 148–154.

[12] D. Heerd, S. Diercks-Horn, M. Fernández-Lahore, Efficient polygalacturonase production from agricultural and agro-industrial residues by solid-state culture of Aspergillus sojae under optimized conditions, SpringerPlus 3(1) (2014) 1–14.

[13] N. Jacob, Pectinolytic enzymes, Biotechnology for Agro-industrial residues utilisation, Springer2009, pp. 383–396.

[14] L. Olsson, T.M. Christensen, K.P. Hansen, E.A. Palmqvist, Influence of the carbon source on production of cellulases, hemicellulases and pectinases by Trichoderma reesei Rut C-30, Enzyme and Microbial Technology 33(5) (2003) 612–619.

[15] E. Käfer, Meiotic and mitotic recombination in Aspergillus and its chromosomal aberrations, Adv Genet 19 (1977) 33–131.

[16] V. Kumar, V. Sahai, V. Bisaria, High-density spore production of Piriformospora indica, a plant growth-promoting endophyte, by optimization of nutritional and cultural parameters, Bioresource technology 102(3) (2011) 3169–3175.

[17] M.M. Bradford, A rapid and sensitive method for the quantitation of microgram quantities of protein utilizing the principle of protein-dye binding, Analytical biochemistry 72(1-2) (1976) 248–254.

[18] G.L. Miller, Use of dinitrosalicylic acid reagent for determination of reducing sugar, Analytical chemistry 31(3) (1959) 426–428.

[19] U.K. Laemmli, Cleavage of structural proteins during the assembly of the head of bacteriophage T4, nature 227(5259) (1970) 680–685.

[20] C.R. Merril, D. Goldman, S.A. Sedman, M.H. Ebert, Ultrasensitive stain for proteins in polyacrylamide gels shows regional variation in cerebrospinal fluid proteins, Science 211(4489) (1981) 1437–1438.

[21] S. Kumar, G. Stecher, M. Li, C. Knyaz, K. Tamura, MEGA X: molecular evolutionary genetics analysis across computing platforms, Molecular biology and evolution 35(6) (2018) 1547.

[22] Y. Chen, D. Sun, Y. Zhou, L. Liu, W. Han, B. Zheng, Z. Wang, Z. Zhang, Cloning, expression and characterization of a novel thermophilic polygalacturonase from Caldicellulosiruptor bescii DSM 6725, International journal of molecular sciences 15(4) (2014) 5717–5729.

[23] S. Verma, A. Varma, K.-H. Rexer, A. Hassel, G. Kost, A. Sarbhoy, P. Bisen, B. Bütehorn, P. Franken, Piriformospora indica, gen. et sp. nov., a new root-colonizing fungus, Mycologia 90(5) (1998) 896–903.

[24] S. Nair, T. Panda, Statistical optimization of medium components for improved synthesis of pectinase by Aspergillus niger, Bioprocess and Biosystems Engineering 16(3) (1997) 169–173.

[25] A.M. Ramos, M. Gally, M.C. García, L. Levin, Pectinolytic enzyme production by Colletotrichum truncatum, causal agent of soybean anthracnose, Revista Iberoamericana de Micología 27(4) (2010) 186–190.

[26] O. Fawole, S. Odunfa, Some factors affecting production of pectic enzymes by Aspergillus niger, International Biodeterioration & Biodegradation 52(4) (2003) 223–227.

[27] G. Aguilar, C. Huitrón, Constitutive exo-pectinase produced by Aspergillus sp. CH-Y-1043 on different carbon source, Biotechnology Letters 12(9) (1990) 655–660.

[28] R.C. Fontana, S. Salvador, M.M.d. Silveira, Influence of pectin and glucose on growth and polygalacturonase production by Aspergillus niger in solid-state cultivation, Journal of Industrial Microbiology and Biotechnology 32(8) (2005) 371–377.

[29] A. Thakur, R. Pahwa, S. Singh, R. Gupta, Production, purification, and characterization of polygalacturonase from Mucor circinelloides ITCC 6025, Enzyme research 2010 (2010) 1–7.

