## Supplementary material for "Polygalacturonase production enhancement by Piriformospora indica from sugar beet pulp under submerged fermentation using surface methodology": graphical abstract

### Optimization

### Characterization

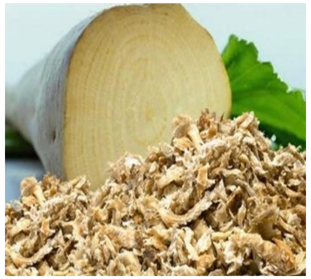

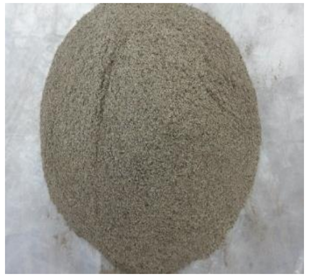

**Sugar beet pulp (SBP)**

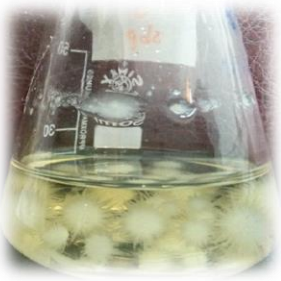

***P. indica***

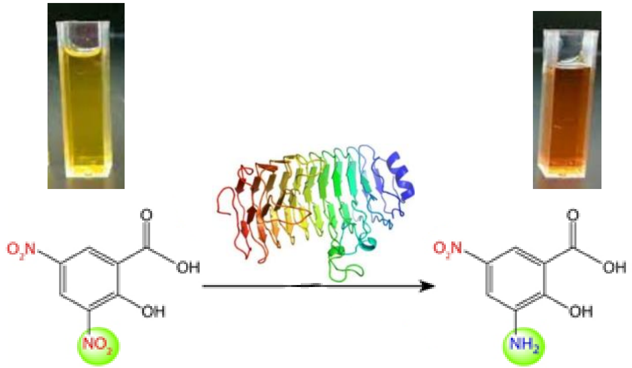


| Run | Variables | | | DW (g/L) | FW (g/L) | pH | Protein (mg/ml) | Observed activity (U/ml) | Specific activity (U/mg) |
|---|---|---|---|---|---|---|---|---|---|
| | A | B | C | | | | | | |
| **1** | 60 | 4 | 20 | 62.56 | 417.85 | 4.09 | 0.25 | **19.41** | 77.64 |
| 2 | 60 | 6 | 15 | 55.5 | 323.36 | 3.72 | 0.18 | 14.74 | 81.88 |
| 3 | 80 | 6 | 15 | 62.06 | 332.6 | 3.55 | 0.157 | 12.33 | 78.53 |
| 4 | 80 | 6 | 10 | 57.53 | 205.86 | 4.7 | 0.151 | 10.46 | 69.27 |
| 5 | 100 | 4 | 20 | 60.85 | 380.85 | 3.78 | 0.196 | 14.16 | 72.24 |
| 6 | 80 | 6 | 15 | 47.77 | 306.98 | 5.09 | 0.148 | 12.13 | 81.95 |
| 7 | 100 | 6 | 15 | 64.39 | 245.88 | 5.76 | 0.135 | 10.76 | 79.70 |
| 8 | 80 | 8 | 20 | 55.94 | 373.6 | 4.85 | 0.433 | 10.12 | 23.37 |
| 9 | 80 | 8 | 15 | 55.335 | 369.56 | 5.06 | 0.217 | 10.67 | 49.17 |
| 10 | 60 | 4 | 10 | 66.20 | 188.61 | 5.29 | 0.145 | 14.03 | 96.75 |
| 11 | 80 | 4 | 15 | 57.11 | 381.42 | 4.28 | 0.148 | 14.03 | 94.79 |
| 12 | 80 | 6 | 20 | 67.33 | 379.35 | 4.01 | 0.186 | 13.28 | 71.39 |
| **13** | 80 | 6 | 15 | 69.22 | 336.53 | 3.88 | 0.105 | 13.33 | **126.95** |
| 14 | 60 | 8 | 20 | 53.35 | 253.26 | 5.60 | 0.252 | 12.35 | 49 |
| 15 | 100 | 8 | 10 | 52.09 | 176.76 | 4.99 | 0.192 | 8.07 | 42.02 |
| 16 | 100 | 4 | 10 | 56.19 | 251.20 | 4.69 | 0.133 | 9.78 | 73.53 |
| 17 | 80 | 6 | 15 | 61.75 | 332.77 | 4.15 | 0.19 | 12.87 | 67.73 |
| 18 | 80 | 6 | 15 | 73.40 | 323.33 | 4.84 | 0.26 | 12.54 | 48.23 |
| 19 | 60 | 8 | 10 | 51.71 | 266.44 | 3.6 | 0.16 | 10.75 | 67.19 |
| 20 | 80 | 6 | 15 | 58.99 | 393.97 | 5.17 | 0.11 | 12.37 | 111.45 |

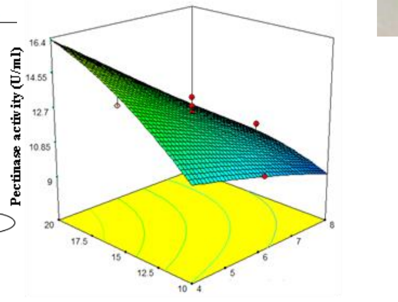


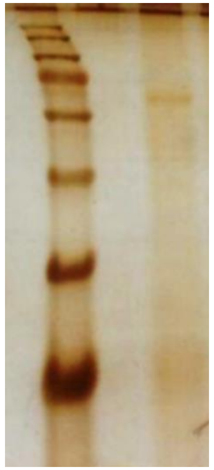

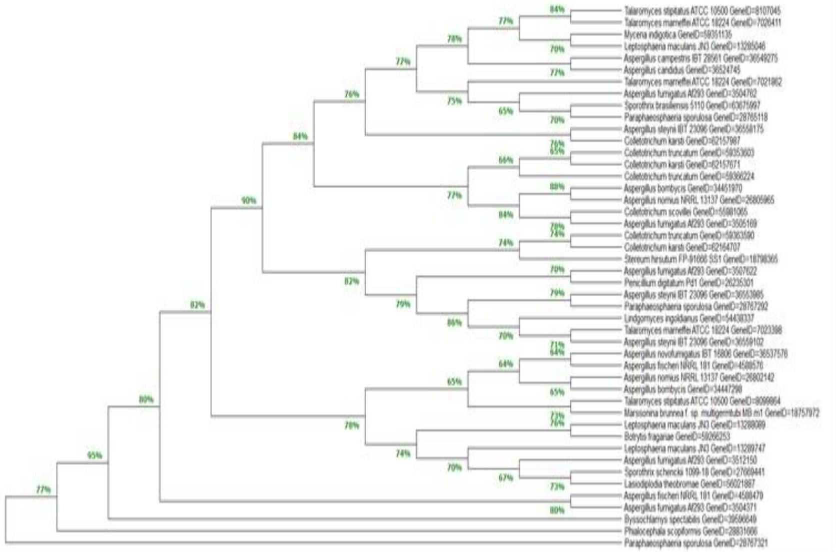

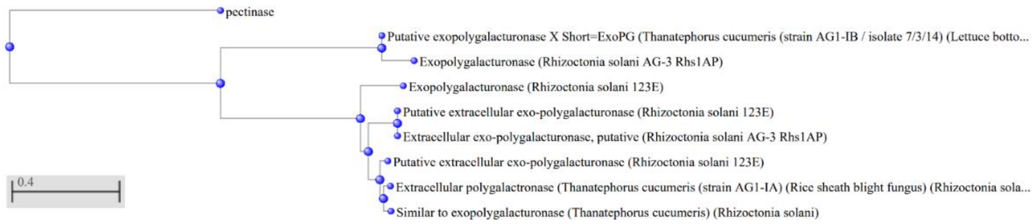
